# SaLTy: a novel *Staphylococcus aureus* Lineage Typer

**DOI:** 10.1101/2023.02.03.527095

**Authors:** Liam Cheney, Michael Payne, Sandeep Kaur, Ruiting Lan

## Abstract

*Staphylococcus aureus* asymptomatically colonises 30% of humans and in 2017 was associated with 20,000 deaths in the USA alone. Dividing *S. aureus* into smaller sub-groups can reveal the emergence of distinct sub-populations with varying potential to cause infections. Despite multiple molecular typing methods categorising such sub-groups, they do not take full advantage of *S. aureus* WGS when describing the fundamental population structure of the species.

In this study, we developed *Staphylococcus aureus* Lineage Typing (SaLTy), which rapidly divides the species into 61 phylogenetically congruent lineages. Alleles of three core genes were identified that uniquely define the 61 lineages and were used for SaLTy typing. SaLTy was validated on 5,000 genomes and 99.12% (4,956/5,000) of isolates were assigned the correct lineage.

We compared SaLTy lineages to previously calculated clonal complexes (CCs) from BIGSdb (n=21,173). SALTy improves on CCs by grouping isolates congruently with phylogenetic structure. SaLTy lineages were further used to describe the carriage of *Staphylococcal* chromosomal cassette containing *mecA* (*SCCmec*) which is carried by methicillin-resistant *S. aureus* (MRSA). Most lineages had isolates lacking SCC*mec* and the four largest lineages varied in SCC*mec* over time. Classifying isolates into SaLTy lineages, which were further SCC*mec* typed, allowed SaLTy to describe high-level MRSA epidemiology

We provide SALTy as a simple typing method that defines phylogenetic lineages (https://github.com/LanLab/SaLTy). SALTy is highly accurate and can quickly analyse large amounts of *S. aureus* WGS. SALTy will aid the characterisation of *S. aureus* populations and the ongoing surveillance of sub-groups that threaten human health.

## Introduction

*Staphylococcus aureus* is a bacterium of major public health concern that asymptomatically colonises ~30% of the human population (1). *S. aureus* causes many forms of disease and is a leading cause of bacteraemia and endocarditis (2). Three waves of *S. aureus* antimicrobial resistance (AMR) have been reported since the 1940s, further complicating the treatment of infections (3). Over this time, *S. aureus* has evolved resistance to multiple routinely used antimicrobials, including virtually all β-lactam antibiotics (methicillin resistant *S. aureus*, MRSA) (4–6). Acquisition of the *Staphylococcal* chromosomal cassette containing *mecA* (SCC*mec*) confers resistance to β-lactam antibiotics. Transmission of *S. aureus*, especially MRSA, was previously linked to healthcare-associated (HA) settings, however, in recent decades community-associated (CA) MRSA in settings with no direct clinical links has been widely reported (7–10).

Separating *S. aureus* into smaller groups can identify the emergence of sub-groups that differ in capacity to cause infection. Numerous molecular typing technologies have been developed to systematically classify these groups (11–13). Pulsed-field gel electrophoresis (PFGE) was once considered the “gold standard” of *S. aureus* classification. PFGE has been widely used to characterise *S. aureus* spread at the continent, country and local levels (14–17). A limitation of PFGE is inconsistency across different laboratories due to variability in interpreting banding patterns (13). The seven-gene multi-locus sequence typing (MLST) method for *S. aureus*, addressed this limitation and enabled standardised comparison between strains (12). Using this approach, the species can be subdivided into sequence types (STs), where isolates with the same ST contain identical alleles at seven loci. MLST has been applied to large collections of isolates and has defined both globally distributed and regionally restricted STs (18, 19). Closely related STs can also be grouped using clonal complexing (20). A clonal complex (CC) contains STs that differ from at least one other ST by a maximum of one allelic difference in the seven genes of the MLST scheme. The genetic similarity of diverse isolates has been described extensively using the CC nomenclature (21–23).

Whole genome sequencing (WGS) been used to generate >80,000 *S. aureus* genomes which can be used to classify the species population structure (24). The extension of MLST to include genes core to a species (core genome MLST, cgMLST) has enabled the high-resolution comparison of thousands of isolates (25–27). In 2014, an *S. aureus* cgMLST scheme was published that consisted of 1,861 core loci (28). cgMLST typing generates an allele profile for each isolate. The allele profile contains the allele calls for each of the 1,861 core loci. The isolates which share the same allele profiles are assigned the same core genome ST (cgST). The high resolution of cgMLST typing is suitable for distinguishing closely related isolates, such as identifying outbreak isolates transmitted within a hospital setting (29, 30). However, cgSTs cannot describe the genetic similarity of diverse isolates since any new allele at any of the core loci will cause the assignment of a novel cgST. The allele profiles generated from cgMLST typing can instead be used to investigate the population structure between diverse isolates using clustering (31, 32). The development of clustering algorithms capable of processing large datasets has facilitated the clustering of cgMLST allele profiles from thousands of isolates (27, 33). Clustering allele profiles has the potential to resolve the fundamental population structure of *S. aureus* by identifying the relationships of highly diverse isolates. To the authors’ knowledge, the clustering of cgMLST allele profiles has not previously been used to resolve the fundamental population structure of *S. aureus*.

In this study, we developed a novel genomic tool for the species-wide classification of *S. aureus* named *S. aureus* Lineage Typer (SaLTy). Developing SaLTy involved (1) dividing the species into lineages through clustering cgMLST allele profiles, (2) selecting alleles of core genes that were specific to the lineages, (3) validating the selected core genes, and (4) designing the SaLTy algorithm. The usefulness of SaLTy was then investigated by dividing over 50,000 *S. aureus* genomes into lineages, showcasing the advantages of lineages over CCs, and characterising the trends in AMR by comparing patterns of AMR within and between SaLTy lineages.

## Methods

### Dataset Curation

*S. aureus* WGS read sets were downloaded from the NCBI Sequence Read Archive (SRA) using NCBI-genome-download (v0.3.1) (34). Contamination was identified with Kraken (v1.0) and read sets with more than 15% of non-*S. aureus* reads were removed (35). Raw reads were assembled by the pipeline developed for the MultiLevel Genome Typer (36). Briefly, the procedure trimmed raw reads with Trimmomatic (v0.39), generated assemblies with Spades (v3.13.1), and calculated assembly quality metrics with Quast (v5.1) (37–39). Assemblies that did not meet the criteria in **Table S1** were removed. STs for the remaining assemblies were called with MLST (v2.10) (40). A species representative dataset was selected with assemblies from each ST. The number of assemblies per ST was proportional to the frequency of that ST. All assemblies assigned a singleton ST were included in the species representative dataset (**Dataset S1**).

### Core Genome Validation

An *S. aureus* core genome was previously defined with 1,861 core loci (28). The core loci of the existing core genome were verified (**Methods S1.1, Dataset S2**). Core genes were validated with isolates from the species representative dataset. The 1,861 loci were called using the Allele Calling Pipeline used in MultiLevel Genome Typing (36). Selected settings were a BLAST (v2.9) similarity of 80% and a 16 SNP sliding window.

### SaLTy Design: species division through hierarchical clustering

Alleles for the species representative dataset were called with the Allele Calling Pipeline used in MultiLevel Genome Typing (36). A local cgMLST database then generated cgMLST profiles by processing the allele calls from each isolate. An in-house python script calculated the pairwise number of allele differences between all allele profiles (**Script 1**). Isolates were separated into clusters using hierarchical clustering of cgMLST with an agglomerative approach (single-linkage clustering). Clusters were defined at a range of allele thresholds (0 to 1,713 alleles). A silhouette index (SI) was calculated for clustering at each allele threshold (**Script 1, Dataset S3**). The phylogenetic relationship of isolates were visualised using GrapeTree (v1.5). GrapeTree generated a neighbour joining phylogeny from allele profiles (41). Isolates of the phylogeny were coloured by their assigned cluster at allele threshold 1,026.

### SaLTy Design: selection of a three gene combination

The definition dataset isolates included half of the representative species dataset. An in-house python script queried the definition dataset to identify alleles of core genes specific to a cluster (**Script 2**). Using a gene-by-gene approach, an allele was assigned to one of the clusters. An allele was assigned to a cluster when present in at least 50% of isolates in that cluster and uniquely carried by isolates of that cluster. The following three performance metrics for every cluster were calculated: accuracy, specificity, and sensitivity (**Methods S1.2**). Each of these metrics was averaged for all clusters to create a macro-accuracy, macro-specificity, and macro-sensitivity. Macro-statistics distributions were visualised in Prism (v9.3.1) (42). Cluster-specific alleles were further selected from two and three gene combinations to increase macro-accuracy where any of the genes could be used to assign a lineage. Single genes with a macro-accuracy in the top 10% were tested in two and three gene combinations. A single three gene combination was selected using a combination of five metrics. The following metrics were used: accuracy, specificity, distance separating genes, number of failed isolates and the isolates in non-redundant lineages (**Methods S1.3**). Each of these metrics progressively filtered three gene combinations until a single combination passed the filtering thresholds in **Table S2**. The alleles from the selected three genes and the clusters they were assigned to type are reported in **Dataset S4**.

### SaLTy Design: validation of the three gene combination

The validation dataset included half of the representative species dataset and was independent of the definition dataset. For each isolate, the alleles for the validated three gene combination were extracted from the previously generated allele profiles. Allele calls for the three gene combination were used to determine the cluster of each isolate by cross referencing with the table of cluster specific alleles in **Dataset S4**. Three gene combination accuracy, sensitivity, and specificity were calculated by comparing the assigned cluster based on allele presence to the cluster defined in hierarchal clustering (**Methods S1.2, Scripts 3 - 4**).

### SaLTy algorithm development and application

The SaLTy pipeline was developed in python 3 (https://github.com/LanLab/SaLTy). Isolates are screened for exact matches using KMA (v1.3.24) (43). The SaLTy pipeline was developed such that it (1) screens isolates against a KMA database of DNA sequences for the cluster specific alleles, (2) filters matches that are 100% in both coverage and identity, and (3) assigns a lineage through cross-referencing the exact allele match with a table of alleles and specific clusters. If an isolate lacks an allele from all three genes, or carries a novel allele for all three genes, it was not assigned a lineage (i.e., was untypable). The SaLTy pipeline was used to analyse the species dataset. All isolates were assigned a SaLTy lineage when a cluster specific allele from any of the three genes was identified. Lineage size distribution was visualised with Prism (v9.3.1) (42). A timing function was built into the SaLTy pipeline for testing multi-threading performance. The seconds required to assign a lineage for each isolate and the total time required to analyse multiple isolates was recorded (**Dataset S6**). The species dataset was analysed in single-threaded and multi-threaded (four CPUs) configurations. For both analyses, 64 megabytes of RAM was allocated. Prism (v9.3.1) visualised the different lineage assignment times between single and multi-threading analysis.

### SaLTy and CC division of *S. aureus* STs from PubMLST

The PubMLST database had 21,883 *S. aureus* genome submissions with associated CC metadata as of April 4, 2020, which were downloaded and typed using SaLTy pipeline. Tableau (v9.1) was used to visualise the grouping of the PubMLST isolates into SaLTy lineages and CCs (44). A subset of 189 isolates from the PubMLST database was selected. The subset represented all CC types and SaLTy lineages. SaLTy lineages with 10 or fewer isolates had all isolates included, and lineages with 10 or more isolates had 10 isolates randomly selected. Snippy (v4.6.0) was used to call SNPs in the subset using GCA_000012045 as a reference (45). SNPs predicted to be affected by recombination were removed with RecDetect (v6.1) (46). A maximum likelihood phylogeny with 10,000 ultrafast bootstraps was generated with IqTree (v2.0.4) (41). Branches of the phylogeny were coloured by bootstrap value, and SaLTy lineage and CC metadata labels were visualised with the Interactive Tool of Life (iTOL) (47).

### *In silico SCCmec* typing

The *SCCmec* type for an isolate was predicted *in silico* with the Staphobia *SCCmec* (https://github.com/staphopia/staphopia-sccmec). Only *SCCmec* types were extracted from prediction results and SCC*mec* subtypes were excluded (**Dataset S7**). *SCCmec* types were reported as ‘Negative’ for isolates that lacked the *mecA* gene. The presence of *mecA* was predicted through Staphobia SCC*mec* analysis. Isolates assigned both *SCCmec* V and VII were typed as ‘SCC*mec* V or VII’. The distribution of SCC*mec* types for isolates of each SaLTy lineage was visualised with Tableau (v9.1) (44).

## Results

### Curation of the species dataset and selection of representative datasets

Developing a *S. aureus* lineage typer required a high-quality genomic dataset. Initially, a species dataset (n=50,481) of quality filtered genomes was created (**Dataset S1**). The species dataset was divided into 1,665 MLST types (**Results S2.1, Figure S1**). A species representative dataset (n=10,000) was selected that sampled each MLST type. The 10,000 isolates in the representative dataset were sampled in proportion to the frequency of each MLST ST in the species dataset. The species representative dataset was further divided into a definition dataset (n=5,000) and a validation dataset (n=5,000). The definition and validation datasets were used to select and validate the core genes used in the SaLTy pipeline.

### *S. aureus* was subdivided into 61 lineages

To subdivide the species, we calculated the allele distance between isolates by comparing cgMLST profiles and then separated isolates into clusters using the allele distances and the hierarchal clustering algorithm. The cgMLST scheme had 1,713 loci present in greater than or equal to 99% of the representative dataset (**Results S2.2, Figure S2**). Allelic profiles for each isolate of the representative dataset were generated from these 1,713 core loci. The allele distance between two isolates was the number of core loci that differed between the allelic profiles of those two isolates. The hierarchal clustering algorithm grouped isolates into clusters at a range of allele thresholds (0 - 1,713). Clusters across the representative dataset were evaluated at each allele threshold using Silhouette Index (SI) (**Figure 1, Dataset S3**). An allele difference of 1,026 was chosen with an SI of 0.815 (3 d.p.). A SI of 0.815 was within the top 1.5% of all SIs and indicated clusters were well separated. An allele difference of 493 had the highest SI (0.824, 3 d.p.), however, we chose an allele threshold of 1,026 as that would cluster isolates into the more genetically divergent lineages and result in more stable clustering (**Figure 1**). Thus, an allele difference of 1,026 alleles was selected that defined 61 clusters. A small number of clusters were found to include a large proportion of isolates, with the five largest clusters (11, 51, 46, 15 and 44) cumulatively assigned to 69.25% (6,925/10,000) of the species representative dataset (**Figure S3A**).

**Figure 1.**
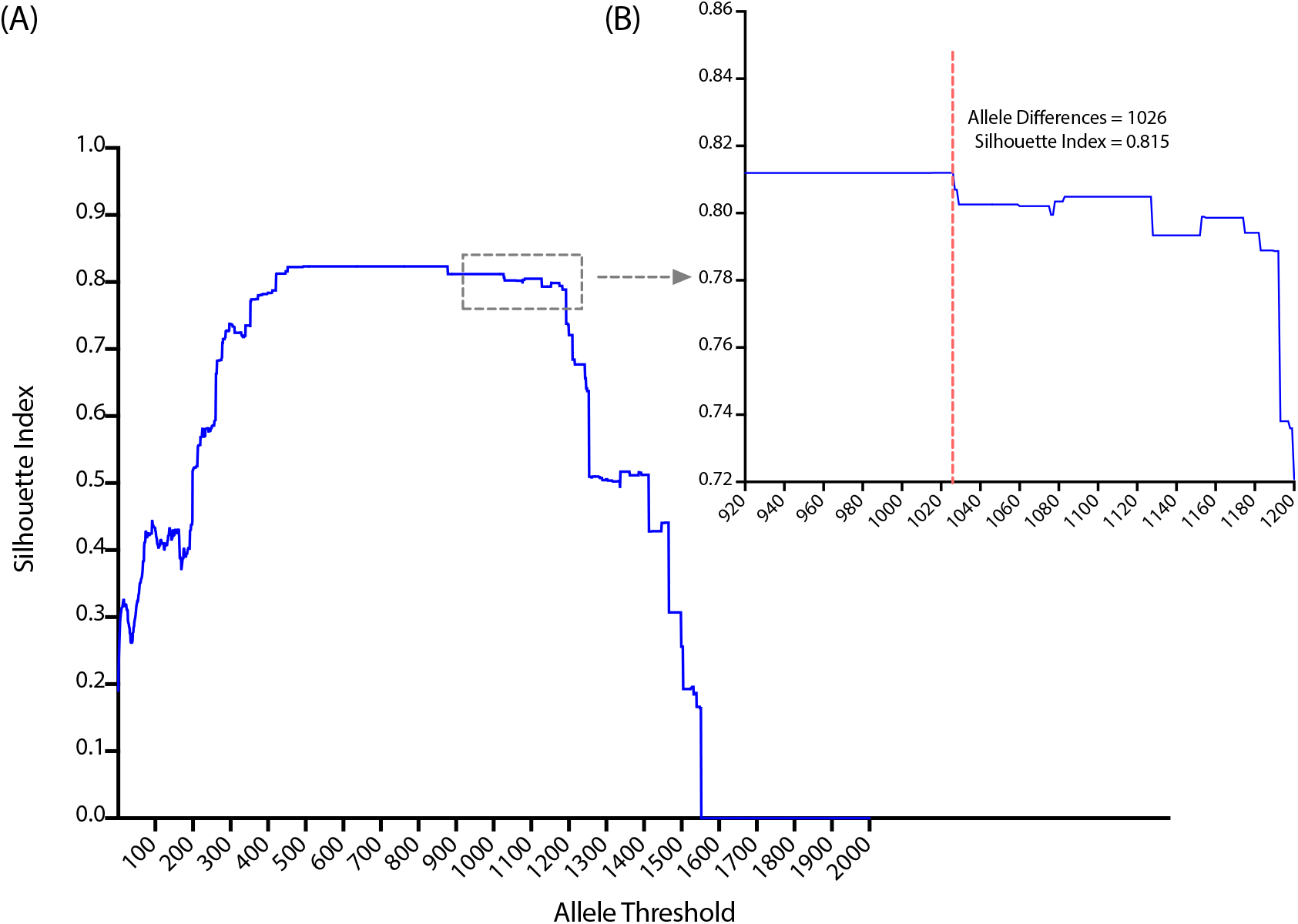
Silhouette indexing supports an allele threshold of 1,026 to divide the species. The species representative dataset (n=10,000) was divided into clusters using hierarchal clustering with cut-offs from zero to 1,713 allelic differences. The Silhouette indexes for each allelic threshold are shown in blue. The dashed line marks the allele threshold selected to cluster the dataset. Visualised in Prism v9.3.1 (42).

Most clusters (49.18%, 30/61) were assigned to less than 20 isolates (Figure S3B). The concordance between the 61 clusters and the species’ phylogenetic structure was investigated (**Figure 2**). Visual inspection showed that all 61 clusters were concordant with lineages of the species phylogeny and therefore were referred to as lineages. In summary, the application of hierarchal clustering defined 61 clusters at an allele threshold of 1,026 with a silhouette index of 0.815. These clusters were referred to as lineages as they were concordant with the phylogenetic structure.

**Figure 2.**
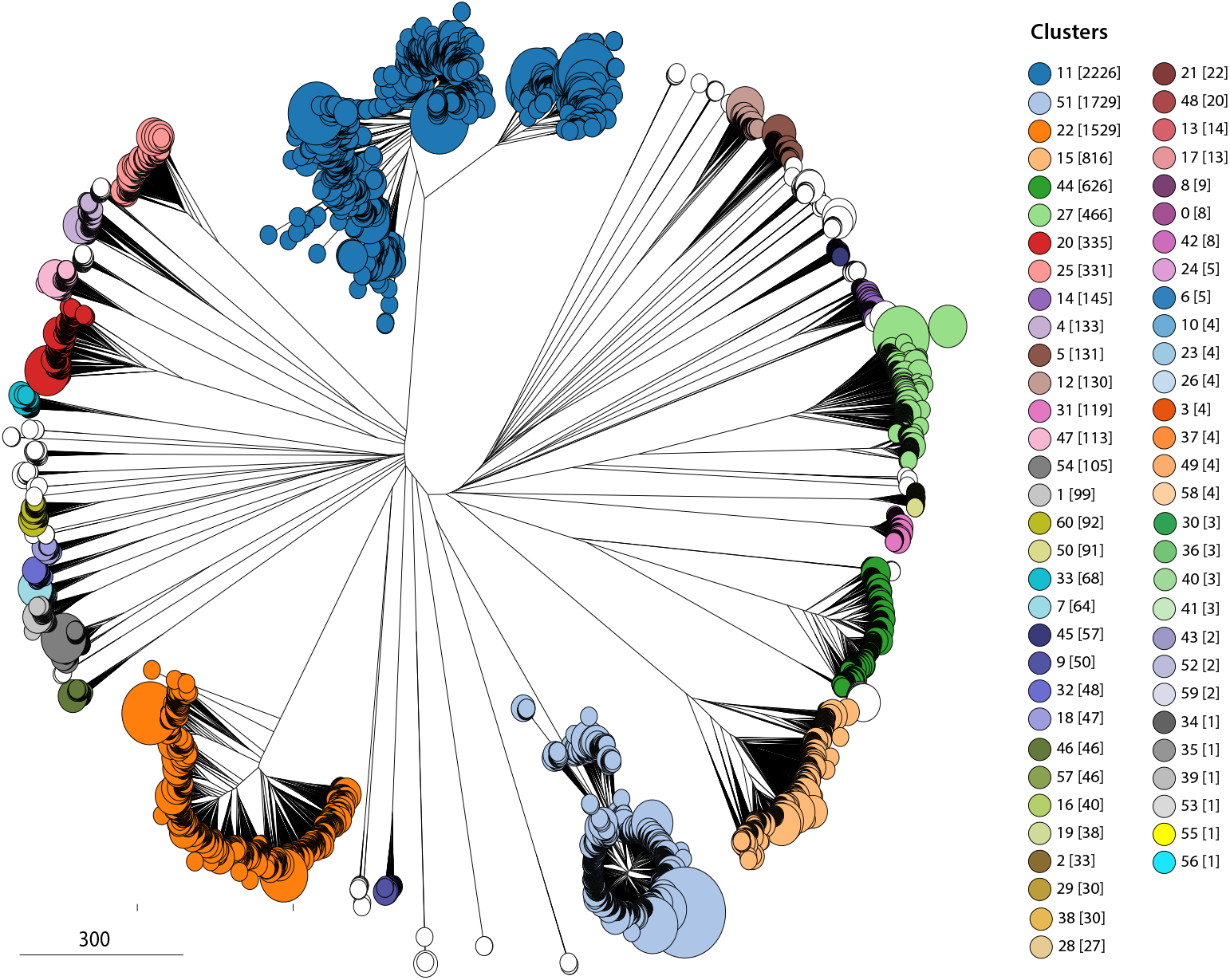
Hierarchal clusters with a 1,026 allele difference cut-off are congruent with the species phylogenetic structure. The phylogenetic structure of the species representative dataset (n=10,000) was compared to 61 hierarchal clusters. Phylogenetic relationships were calculated from cgMLST profiles using the Rapid Neighbour Joining algorithm. Isolates were coloured by cluster type. The number of isolates assigned to each cluster was shown in square brackets. Clusters with less than or equal to 45 isolates were coloured white. The phylogeny was visualised in GrapeTree (41).

### Selection of three genes with alleles specific to the 61 Lineages

Determining the lineage for a new isolate that was not included in the initial analysis would have required re-computing a distance matrix and repeating the hierarchal clustering and thus would be difficult and time consuming. To solve this problem, we aimed to identify individual core genes with alleles specific to as many of the 61 defined lineages as possible (**Results S2.3, Figure 3, Dataset S4**). We examined single genes and combinations of two and three genes (**Figure 3, shown in red and green, Dataset S4**). A combination of genes allowed a lineage to be assigned using an allele from one or more of the genes in a combination. Assigning a lineage did not require alleles for all genes in a combination. The accuracy of lineage assignment using a single gene, and two or three gene combinations was compared (**Figure 3, marked by asterisk, Dataset S4**). A gene combination with a higher accuracy had alleles that grouped isolates more similarly to the 61 defined lineages. As the number of genes increased in a combination, so did the accuracy of the best performing combination. Alleles from single, two and three gene combinations had maximum accuracies of 87.94%, 97.90% and 99.78% respectively (**Figure 3, marked by asterisks**). Thus, three gene combinations provided higher accuracy. A single three-gene combination was selected using a combination of five metrics (**Table S3, Dataset S4**). The selected three gene combination was SACOL0451-SACOL1908-SACOL2725. For the five metrics, this combination had an accuracy of 99.71%, a specificity of 100%, the distance between the genes of at least 100,000 base pairs, an allele for assigning lineages in 99.2% of the definition dataset, and lineage assignments made without any redundancy (called by only 1 out of 3 genes) only occurred in 1.93% of the definition dataset.

**Figure 3.**
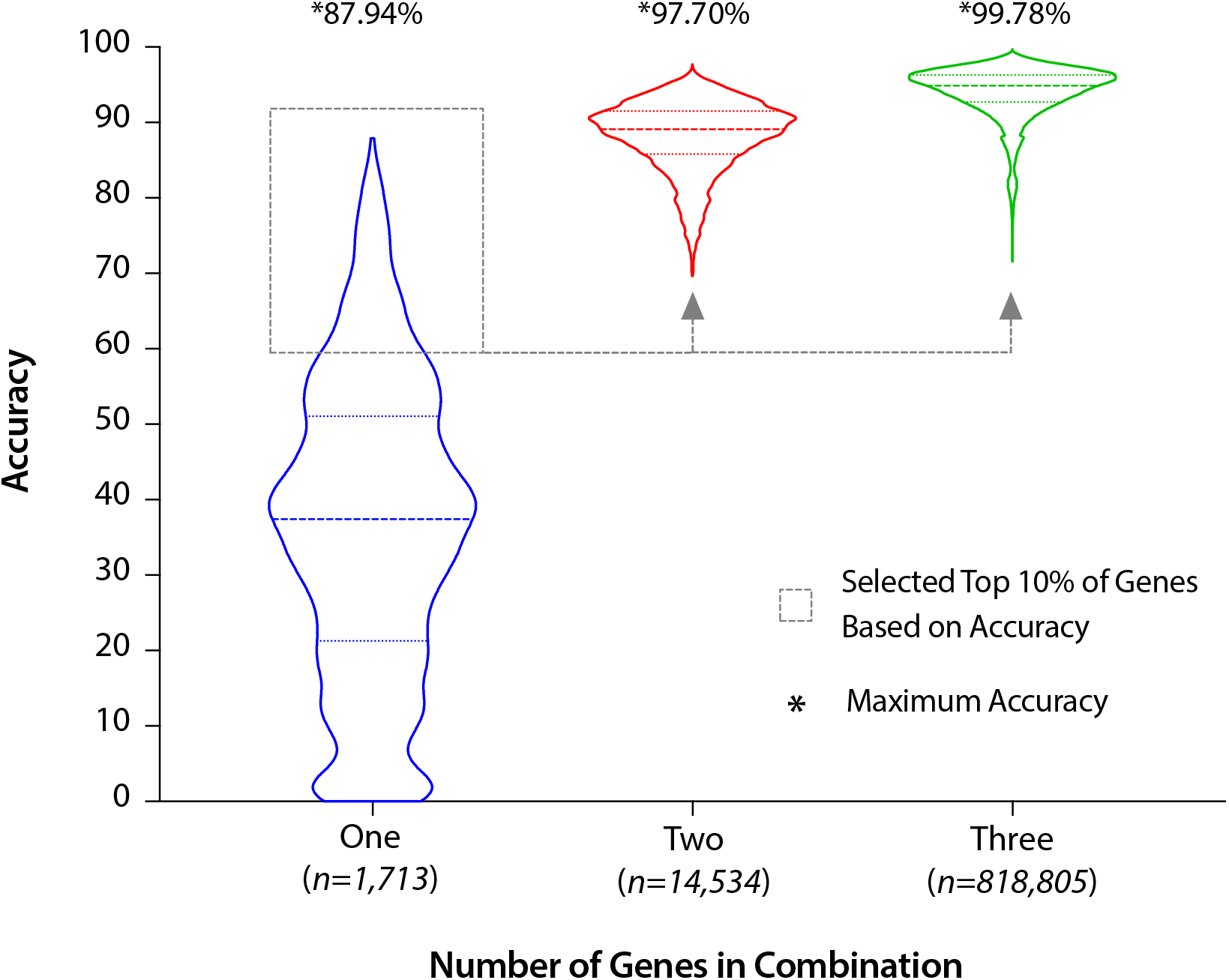
Increasing the number of genes used for specific alleles increased the accuracy of assignment to the 61 hierarchal clusters. Accuracy of one, two and three core gene combination in grouping isolates into the 61 hierarchal clusters was visualised. Initially, alleles were identified that were specific to more than 50% of isolates of a hierarchal cluster on a gene-by-gene basis. The accuracy for each core gene was shown in blue. Based on the accuracy, the top performing 10% individual genes were selected to create two and three combinations (shown by grey dashed line). The number of two and three gene combinations was shown below the X-axis. The accuracy for two and three gene combinations was calculated by testing for the presence of an allele that is specific for a cluster in any of the genes in a combination. The accuracy for two and three gene combinations was shown in red and green (respectively). The maximum accuracy for one, two and three gene combinations was labelled and marked by an asterisk.

### Independent validation of lineage typing using a three gene combination

To validate the selection of the three genes, we assigned SaLTy lineages to isolates of a validation dataset *(n=5,000)*. These isolates were not included when selecting a three gene combination. However, they were a part of the original hierarchal clustering. SaLTy was used to screen isolates for a lineage specific allele from at least one of three core genes and was able to assign a lineage to 99.56% (4,978/5,000) of the validation dataset (**Dataset S4**). All lineage assignments were correct (**Dataset S4**).

### *S. aureus* population structure as described by SaLTy lineages

The three validated genes, SACOL0451-SACOL1908-SACOL2725, were the basis of SaLTy typing. The alleles of these core genes acted as markers for determining a lineage without hierarchal clustering. The SaLTy pipeline analysed an isolate by (1) reporting the alleles for each of the three core genes (2) assigning a lineage by cross referencing the reported alleles with a table of alleles specific to the 61 lineages. SaLTy was used to divide the species dataset (n=50,481) into lineages (**Dataset S5**). SaLTy successfully assigned a lineage to 99.25% (50,100/50,481) of isolates (**Figure 4**). The three largest lineages were 11, 51 and 46, which included 54% (24,335/50,481) of the genomes (**Figure 4A**). There were seven lineages with 10 or fewer isolates, and lineage 55 was the only singleton (**Figure 4B**). The time required to assign an isolate to a lineage and the total time required to analyse the species dataset was recorded **(Dataset S6)**. On average, an isolate was assigned a SaLTy lineage in 0.133 seconds, and the entire species dataset was analysed in 112 minutes and 3 seconds (**Figure S4, shown in red**). The initial analysis of the species was a single-threading application (single CPU). To investigate the multi-threading capability of the SaLTy pipeline, the species dataset was re-analysed with four threads. SaLTy multi-threading analysed the species dataset five times quicker and assigning the 50,481 isolates to a lineage took 24 minutes and 55 seconds (**Figure S4, shown in blue**).

**Figure 4.**
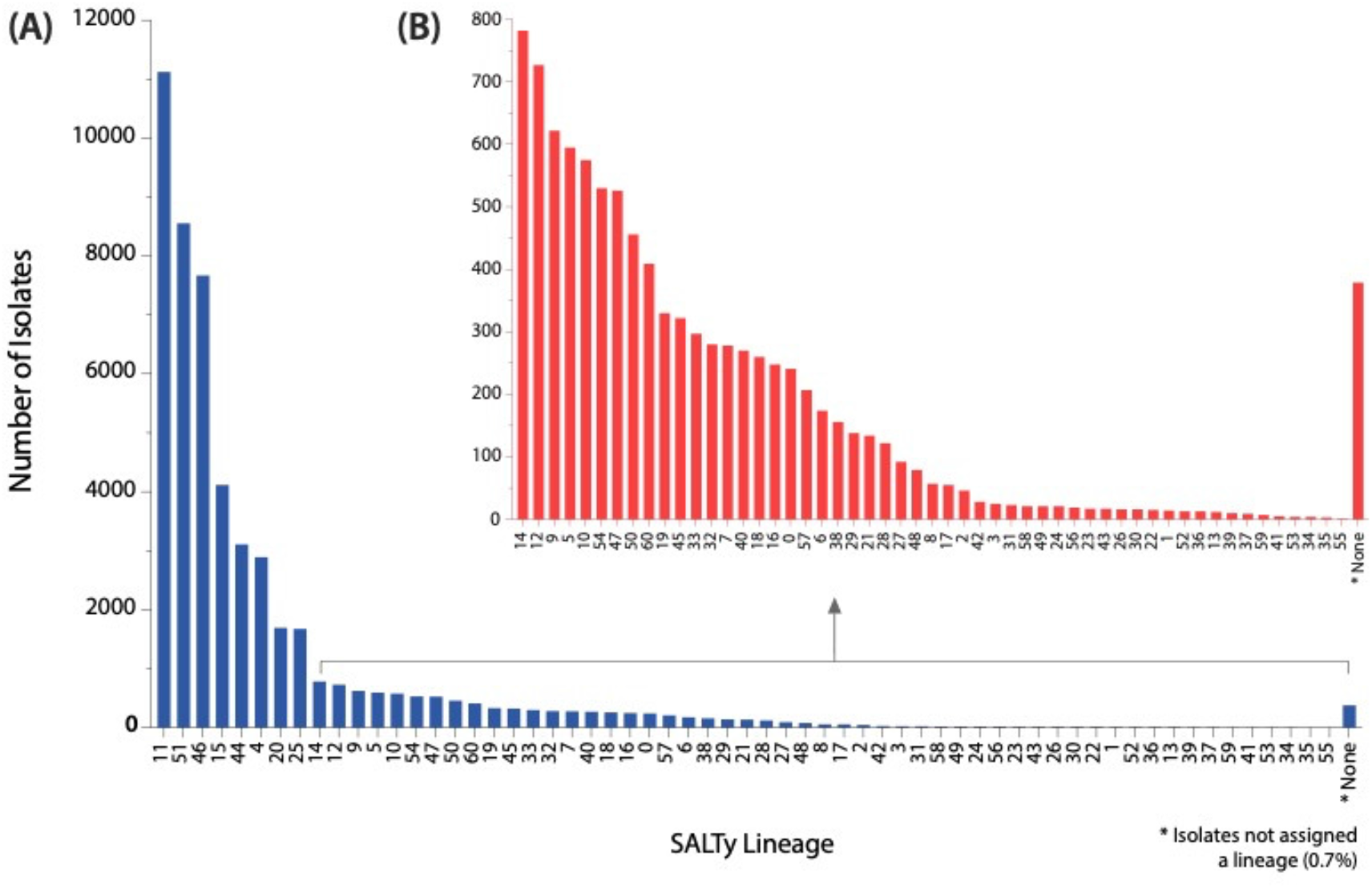
SaLTy divided the *S. aureus* species into 61 lineages. SaLTy processed and assigned a lineage to 99.3% of the complete species dataset (n=50,481). Isolates not assigned a lineage were represented in the “None” category (marked by asterisk). (A) The distribution of the sizes of all lineages across the dataset. (B) A subplot of cluster sizes that were assigned to less than 800 isolates. Visualised in Prism v9.3.1 (42).

### SaLTy lineage typing has a higher resolution than Clonal Complexing and SaLTy lineages are congruent with the Phylogeny

The concordance of the *S. aureus* clonal complexes and SaLTy was investigated. The PubMLST database contained 21,173 *S. aureus* genomes that were assigned CCs and were classified into 10 CCs (48). Using SaLTy, these isolates were divided into 21 lineages. Comparing the SaLTy lineages to CCs showed that 50% (5/10) of CCs were split into multiple SaLTy lineages (**Figure S5**). SaLTy split CC1, CC5, CC8, CC97 and CC121, into three or more lineages each. Isolates in the remaining CCs, CC15, CC22, CC30, CC45 and CC93, were respectively typed by lineages 25, 51, 15, 4 and 47.

The groups of isolates defined by SaLTy and CC were compared by overlaying their assignments onto a maximum likelihood phylogeny (**Figure 5**). The phylogeny included representatives from each of the 21 SaLTy lineages and 10 CCs. Annotating SaLTy lineages alongside the phylogeny showed SaLTy grouped isolates congruently with clades of the phylogeny. All SaLTy lineages (21/21) included isolates from single phylogenetic clades. In contrast, Clonal Complexing defined CCs that combined multiple phylogenetic clades (**Figure 5, highlighted in yellow, green, red, blue and grey**). Half of the CCs (5/10) each combined multiple clades of the phylogeny. CC1, CC5, CC8, CC97 and CC121 all included two or more phylogenetic clades that SaLTy separated into unique lineages. For example, CC1 included isolates from four clades (**Figure 5, shown in yellow**). SaLTy divided these isolates into three lineages 5, 12 and 24 that each formed a distinct clade.

**Figure 5.**
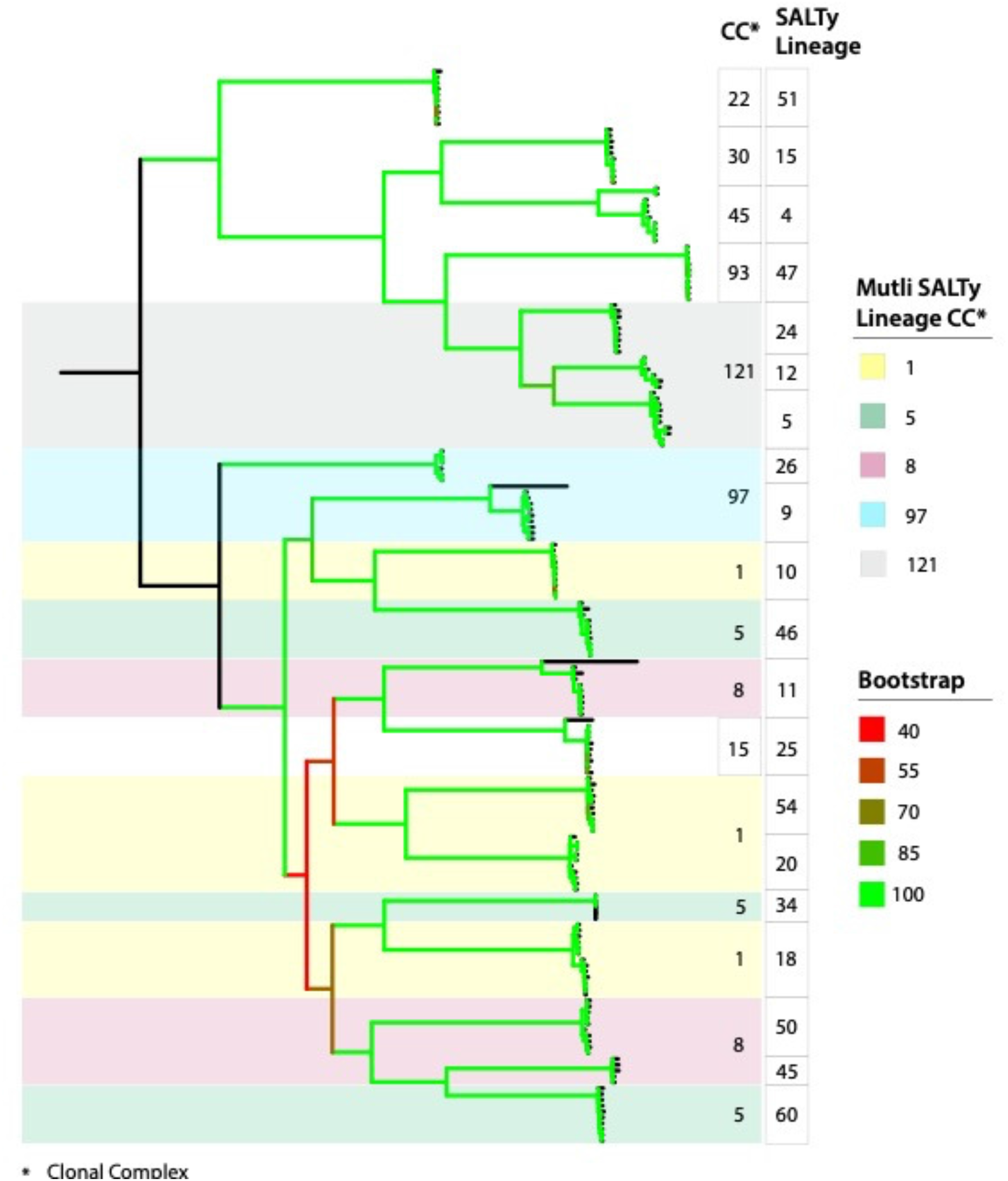
SaLTy splits clonal complexes (CCs) into multiple lineages that are congruent with the phylogeny. A maximum-likelihood phylogeny of *S. aureus* isolates (n=185) from multiple SaLTy lineages and CCs. The phylogeny was based on 33,275 non-recombinant SNPs. SaLTy lineages and CCs were annotated alongside. CCs split into multiple SaLTy lineages were coloured (shown in yellow, green, red, blue and grey). Branches were coloured by ultra-fast bootstrap values (n=10,000). Phylogeny was generated using IQ-TREE v2.0.4 and visualised in iTOL (47, 80).

### SaLTy lineages are heterogeneous in *SCCmec* type and the largest SaLTy lineages vary in *SCCmec* type over time

To describe the grouping of MRSA and MSSA isolates, we predicted the *SCCmec* type for isolates in each SaLTy lineage (n=50,100). Isolates with a predicted SCC*mec* element were assumed to be MRSA, and isolates without a predicted *SCCmec* were assumed to be MSSA. Most lineages (59.02%, 36/61) had at least one isolate predicted to carry *SCCmec* (**Dataset S7**). Five different *SCCmec* types (SCC*mec* I, II, III, IV and V or VII) were predicted in the 50,100 isolates in these 36 SaLTy lineages (**Figure S6**). There were 14 lineages with two or more *SCCmec* types and 12 lineages with only a single SCC*mec* type. The lineages with multiple SCC*mec* types were considerably larger than lineages with a single SCC*mec* type. The largest lineage with multiple SCC*mec* types was lineage 11, which included 11,119 isolates, and the largest lineage with a single SCC*mec* type was lineage 5, which included 590 isolates (**Figure S6A**). There were 34 remaining lineages had no isolates with a predicted *SCCmec* type (**Figure S6, shown in blue**). Cumulatively, these *SCCmec* negative lineages contained 4,543 isolates and were generally smaller than lineages with isolates carrying a SCC*mec* element. The largest lineage without any isolates carrying an *SCCmec* element was lineage 25 which had 1,671 isolates (**Figure S6A**).

The changing frequencies of SCC*mec* types over time from 2001 to 2019 in the four largest SaLTy lineages were visualised (**Figure 6**). Lineages 11, 15 and 46 all had a mixture of SCC*mec* types in almost all years sampled. Lineage 11 had both *SCCmec* III and IV between 2000 to 2008, and from 2009 onwards, the majority of isolates were *SCCmec* IV (**Figure 6A, shown in green)**. Lineage 15 was predominately *SCCmec* II before 2012, but post-2012, was a combination of *SCCmec* II and IV (**Figure 6B**). Additionally, lineage 15 was the only lineage to continually increase in the number of SCC*mec* negative isolates over time. Lineage 46 was a mixture of *SCCmec* II and IV each year, and within that mixture *SCCmec* II was more common for all years (except 2019) (**Figure 6C, shown in red**). Finally, lineage 51 was almost homogeneous in *SCCmec* type with *SCCmec* IV being the only type in almost all years (**Figure 6D, shown in green**). The only other SCC*mec* type in lineage 51 was *SCCmec* V or VII that was found in a very small number of isolates (**Figure 6D, shown in grey**).

**Figure 6.**
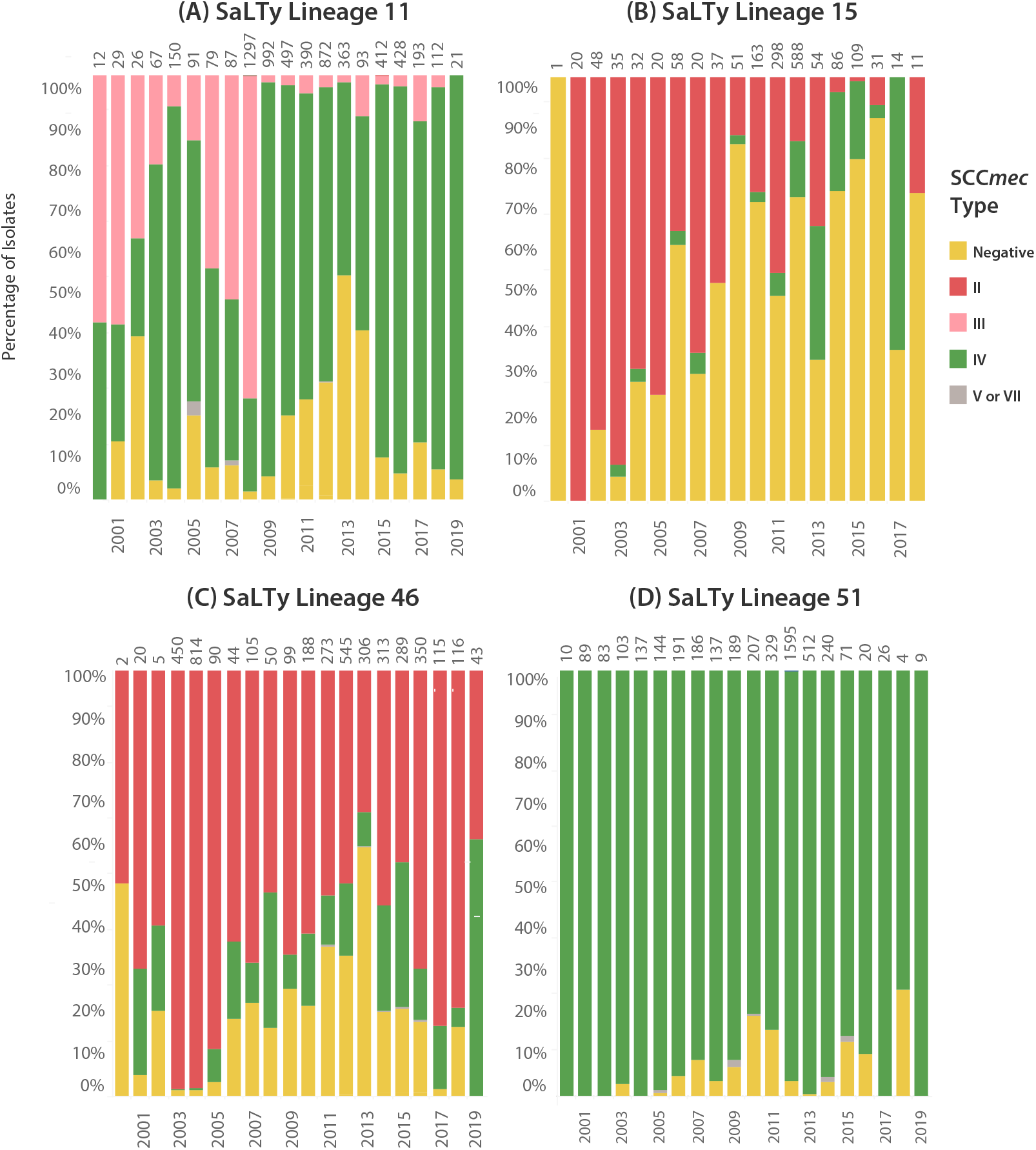
Comparison of the *SCCmec* types over time in SaLTy lineages 11, 15, 46 and 51. The SCC*mec* types in SaLTy lineages 11, 15, 46 and 51 between 2000 to 2019 were compared. *SCCmec* negative isolates were yellow, and *SCCmec* II, III, IV and V and VII were coloured as red, pink, green and grey (respectively). The frequencies of predicted SCC*mec* types were shown as the percentage of the total isolates per year. The total number of isolates sampled from each year was shown above each column.

## Discussion

### SaLTy offers rapid whole species classification

WGS has ushered in a new era of *S. aureus* genomic epidemiology. The analysis of *S. aureus* WGS data is growing in popularity and will likely become routine for species-wide surveillance (49, 50). The majority of *S. aureus* WGS data has been generated in the past decade, which offers a new opportunity to develop rapid and scalable WGS-based classification technologies (51). In this study, we developed a novel genomic typing technology named SaLTy. SaLTy is a rapid genome typer that divides the *S. aureus* species into groups consistent with phylogenetic lineages. The typing resolution of SaLTy is lower than traditional MLST and comparable to that of Clonal Complexing (**Figure 5**) (28, 52). Unlike MLST and clonal complexing, the SaLTy lineages were defined through the high-resolution investigation of thousands of cgMLST allele profiles, and the lineages represent the fundamental population structure of the species (**Figure 2**).

Currently, there is a lack of genomic classification technologies that can rapidly assign lineages using *S. aureus* WGS. The large-scale population structure is most often resolved through phylogenetic reconstruction (53–55). Databases have been developed that can build phylogenies with tens of thousands of isolates and generate interactive visualisations that aid in investigating genomic epidemiology (27, 41). However, incorporating newly sequenced *S. aureus* WGS data into a phylogeny requires rerunning the entire analysis. This makes phylogenetic reconstruction unfeasible for the ongoing interpretation of newly sequenced *S. aureus* isolates. SaLTy can assign a lineage by screening isolates for alleles from three pre-selected core genes. SaLTy has been used to process over 50,000 *S. aureus* genomes and can be used to process future *S. aureus* WGS data (**Figure 3**). We provide SaLTy as a freely available genomic tool (https://github.com/LanLab/SaLTy) to assign lineages that indicate the genetic diversity of *S. aureus* and facilitate the ongoing monitoring of existing and newly emerging *S. aureus* lineages (56, 57).

### SaLTy lineages describe MRSA and MSSA epidemiology

*S. aureus* is notorious for its ability to acquire genetic elements that confer resistance to antibiotics (58–61). Traditional typing technologies have defined multiple sub-groups that have acquired AMR elements and proceeded to expand globally (62–65). SaLTy separated the species into lineages of diverse isolates and found that over half the lineages (36/61) contained MRSA isolates (**Figure 6, Figure S6**). The genetic basis of MRSA is the presence of *SCCmec* which can be transmitted between isolates through mechanisms of HGT. Variants of *SCCmec* have also been characterised that contribute to varying MRSA resistance profiles (66). In one landmark study, the emergence of novel SCC*mec* types was used to define multiple waves of MRSA endemic spread (3). We compared the changing frequencies of SCC*mec* types over time and showed the four largest SaLTy lineages varied in the carriage of SCC*mec* type. By describing the trends in SCC*mec* types in SaLTy lineages, we could show that large SaLTy lineages did not always carry the same SCC*mec* types, which is relevant when tracking common antibiotic resistance phenotypes.

In the study of the evolution of MRSA, the classification of isolates lacking *SCCmec* (MSSA) must also be considered. There were 34 lineages with all MSSA isolates. Classifying lineages of MSSA is important as MSSA isolates can acquire the *SCCmec* and convert into MRSA (67). This phenomenon is not rare and the emergence of MRSA in the species has been shown to have independently occurred multiple times (68, 69). The acquisition of *SCCmec* can lead to rapid expansion of a lineage and SaLTy will be able to identify such lineages rapidly and accurately.

MSSA lineages that remain SCC*mec*-free are also essential to classify. MSSA are the largest cause of *S. aureus* related disease(s) and recent reports have found a resurgence in MSSA infections and a reduction in MRSA infections since the 2000s (70–73). The ability for SaLTy to rapidly type new isolates, which can further be SCC*mec* typed (SCC*mec* positive or SCC*mec* negative) allows SaLTy to describe high-level MRSA epidemiology. These lineages can spread within hospital or community settings and complicate disease treatments. By characterising the lineages of MRSA with SaLTy, the spread of existing or newly emerging MRSA can be described.

### SaLTy lineages represent the basic population structure of the species

An essential aspect of the genomic epidemiology *of S. aureus* is resolving its population structure. A detailed understanding of the large-scale population structure of the species is the basis for investigating the evolution of both well-established and emerging sub-groups (74, 75). Large-scale population structures have been resolved for other species by clustering thousands of allele profiles generated with cgMLST typing (27, 75, 76). Resolving the species-level population structure of *S. aureus* through clustering cgMLST allele profiles would require selecting an allele threshold that grouped genetically diverse *S. aureus*. The SaLTy lineages were defined by separating the *S. aureus* species into 61 lineages that differed by no more than 1,026 core genes (**Figure 2, Figure 4**). The 1,026 allele threshold was selected by adopting a previously described method of silhouette index evaluation (75). The 61 lineages are expected to group distantly related isolates and represent ancestral stages of evolution (**Figure 2**). Even though the likelihood of isolates not being a part of the 61 lineages increases as sequencing continues, an advantage of the approach used in SaLTy is that new alleles and lineages can be defined while maintaining existing definitions.

The current description of *S. aureus* fundamental population structure has been achieved with Clonal Complexing (22, 52). In 2003, a snapshot of the species was generated with the Clonal Complexing of 334 *S. aureus* isolates (22). Clonal Complexing identified CCs that represented most of the species and CCs associated with global MRSA transmission. We compared Clonal Complexing and SaLTy division of the species and showed that only half of the CCs (5/10) were assigned a single SaLTy lineage (**Figure 5**). Further investigation of the remaining CC showed SaLTy divided these CCs into lineages consistent with the underlying phylogenetic signal. A limitation of Clonal Complexing is that CCs based on alleles from the seven gene MLST scheme do not reflect phylogenetic structure (20). SaLTy lineages instead grouped isolates congruently with the phylogenetic structure, which is theoretically a true representation of the species population structure. The SaLTy lineages provide an up-to-date representation of the *S. aureus* species-level population structure that was generated through the detailed analysis of a large volume of *S. aureus* WGS data. Resolving the species-level population structure with SaLTy lineages will aid the future description of *S. aureus* large-scale epidemiology.

### The potential extension of SaLTy in a PCR assay

Although SaLTy was initially developed to process *S. aureus* WGS data and assign a lineage, SaLTy could be developed into a PCR-based amplicon typing technology (77–79). The alleles from the three SaLTy genes could be used through (1) designing three simplex PCR assays or a single triplex PCR assay, (2) amplicon sequencing the PCR products, and (3) assigning a lineage through processing the amplicon sequencing with existing SaLTy genomics pipeline. SaLTy as a PCR-based typing technology could offer high-level division of the species that current classification methods cannot. The cost of WGS could act as a barrier to the adoption of SaLTy. Developing SaLTy into a laboratory-based method with amplicon sequencing that can be highly multiplexed would facilitate faster and larger scale lineage typing of *S. aureus* isolates without the need for WGS.

## Conclusion

*S. aureus* is a significant burden to public healthcare systems globally and is the leading cause of hospital acquired infections. Recent advances in WGS technologies have led to the sequencing of tens of thousands of *S. aureus* isolates. There is a new opportunity to describe *S. aureus* genomic epidemiology through the development of scalable WGS-based typing technologies.

In this study, we developed a novel genomic typing technology named SaLTy. SaLTy assigns a lineage to *S. aureus* WGS data and is suitable for processing large volumes of WGS. SaLTy was used to describe the high-level population structure of the species and resolved the species into 61 lineages. SaLTy lineages were compared to Clonal Complexing and showed that lineages improved on CCs by grouping isolates congruently with phylogenetic structure. SaLTy lineages were further used to describe the carriage of SCC*mec* throughout the species. We showed that (1) the majority of SaLTy lineages were MSSA, and (2), the four largest SaLTy lineages carried different frequencies of *SCCmec* variants over time.

We provide SaLTy as a novel WGS-based typing tool for describing large-scale *S. aureus* genomic epidemiology. The SaLTy pipeline and the lineages assigned to the species are freely available (https://github.com/LanLab/SaLTy). We anticipate SALTy will facilitate the identification and ongoing surveillance of existing and emerging *S. aureus* lineages that poses a threat to human health.

## Acknowledgments

The authors would like to thank Duncan Smith and Robin Heron from the UNSW Restech Technology Services Team for their ongoing assistance with high powered computing and data management systems.

